# Network Inference Reveals Distinct Transcriptional Regulation in Barley against Drought and Fusarium Head Blight

**DOI:** 10.64898/2025.12.09.693163

**Authors:** Christina E. Steidele, Johannes Kersting, Felix Hoheneder, Markus List, Ralph Hückelhoven

**Affiliations:** Chair of Phytopathology, TUM School of Life Sciences, HEF World Agricultural Systems Center, Technical University of Munich, Emil-Ramann Str. 2, 85354 Freising-Weihenstephan, Germany; Data Science in Systems Biology, TUM School of Life Sciences, Technical University of Munich, Maximus-von-Imhof-Forum 3, 85354 Freising-Weihenstephan, Germany; Munich Data Science Institute (MDSI), Technical University of Munich, Walther-von-Dyck-Straße 10, 85748 Garching, Germany

**Keywords:** Barley, *Hordeum vulgare*, Fusarium head blight, gene regulatory network, GENIE3, weighted gene co-expression network analysis, WGCNA, transcription factor binding sites, Paired motif enrichment tool, PMET

## Abstract

We analyzed transcriptional networks in barley under single and combined Fusarium head blight (FHB) and drought stress. We applied complementary Weighted Gene Correlation Network Analysis (WGCNA) to identify stress-associated gene co-expression modules and GENIE3 to infer gene regulatory networks (GRNs). Integration of these frameworks revealed strong overlaps between co-expression modules and GRN clusters, highlighting robust regulatory patterns. Key transcription factors (TFs) were identified based on their weighted node degrees, reflecting their connectivity within the network. Independent analysis of paired transcription factor binding sites in promoter regions further supported predicted regulatory interactions. Notably, WRKY TFs emerged as central regulators of FHB response, consistent with their known roles in defense and secondary metabolite biosynthesis, but did not appear in drought-associated contexts. For bHLH or NAC TFs, individual family members steered FHB or drought responses but not both. Our findings demonstrate the power of combining network inference and motif enrichment to identify candidate TFs controlling stress responses, providing a solid foundation for targeted functional validation.

## Introduction

Barley is the fourth most important cereal crop globally. Like any plant, it faces various abiotic and biotic stresses, and malting barley production is endangered by climate change-related drought and heat (Xie et al., 2018). A recent study has predicted that diseases caused by fungal soil-borne pathogens will increase due to a warming climate (Delgado-Baquerizo et al., 2020). This trend could make agriculture more vulnerable, coinciding with a projected rise in global food demand. Consequently, this situation may result in widening gaps in food supply. Understanding plant physiology in the context of complex stress combinations may help in breeding crops that are resistant to diseases and tolerant to abiotic stress (Pandey and Senthil-Kumar, 2019). Fungi from the Fusarium genus are soil-borne pathogens that affect small-grain cereals such as barley, wheat, rye, and oats, causing Fusarium head blight (FHB), particularly *Fusarium graminearum*, but also *Fusarium culmorum, Fusarium avenaceum* or others. Head infection occurs during the flowering stage, especially under warm and humid conditions (Parry et al., 1995), resulting in discolored and shriveled kernels. A significant concern associated with FHB is the contamination of grains with mycotoxins, particularly trichothecenes such as deoxynivalenol (DON), which pose health risks to both humans and livestock animals. DON is a potent protein biosynthesis inhibitor in eukaryotic cells, exerting its toxic effects by binding to the peptidyl transferase center of the 60S ribosomal subunit, thereby disrupting translation elongation. Exposure to DON results in symptoms such as vomiting (hence its traditional name “vomitoxin”), reduced feed intake, immunosuppression, and gastrointestinal disturbances (Foroud et al., 2017). Complete resistance to FHB in barley or other grasses is absent. FHB resistance appears to be a quantitative trait influenced by multiple genomic loci and highly depends on genotype-environment interactions (Miedaner et al., 2024). Unlike complete resistance, partial resistance in barley does not entirely prevent infection but significantly reduces disease severity and DON accumulation in the kernels. This form of resistance is typically governed by quantitative disease resistance (QDR), which involves multiple quantitative trait loci (QTLs) contributing incrementally to the plant’s defense mechanisms (Buerstmayr and Lemmens, 2015). Significant QTLs have been identified on barley chromosomes 2H, 3H, 5H, 6H, and 7H (Ogrodowicz et al., 2020; Huang et al., 2021). However, the effectiveness and stability of these QTLs can vary depending on the genetic background of the barley cultivar and environmental conditions, highlighting the complexity of breeding for FHB resistance (Linkmeyer et al., 2013; Huang et al., 2021; Sallam et al., 2024). Drought can enhance or mitigate FHB resistance of barley depending on the sequence of environmental challenges and host genotype (Hoheneder et al., 2023, 2025). It is therefore important to study regulatory mechanisms that underlie drought and FHB responses to gain a better understanding of genotype-environment interactions in the future. Transcription factors (TFs) are proteins that regulate gene expression by binding to specific DNA sequences, influencing the recruitment of RNA polymerase. They recognize short DNA motifs, often located within promoter or enhancer regions, and they work in coordination with other TFs or regulatory proteins to fine-tune transcriptional activity. Their roles are essential for responding to developmental cues, environmental signals, and stress conditions. Alterations in the expression of TFs have been shown to influence disease resistance in crops. For instance, overexpression of two barley WRKY domain-containing transcription factors (WRKY) TFs (HvWRKY6 and HvWRKY70) in wheat improved the resistance to stripe rust (*Puccinia striiformis f. sp. tritici*) and powdery mildew (*Blumeria graminis f. sp. tritici*) Li et al. (2020) and overexpression of a wheat NAM/ATAF/CUC (NAC) TF TaNACL-D1 enhances resistance against *F. graminearum* (Vranić et al., 2023). Knockdown of HvWIN1 (Ethylene Response Factor/ APETALA2, ERF/AP2 TF) increased disease severity of FHB in barley (Kumar et al., 2016).

In a previous study, Hoheneder et al. (2023) performed 3’RNA-sequencing on four barley cultivars subjected to drought stress, artificial inoculation with *F. culmorum* spore solution, or a combination of both. Their results demonstrated that barley exhibits a modular response to these combined stresses, suggesting an additive composition of the various stress responses rather than a specific response to the combination of stresses.

This study aims to identify key TFs that act as central regulators of both FHB and drought responses, along with their potential target genes, using an existing transcriptomic dataset from Hoheneder et al. (2023). To achieve this, we constructed a GRN with the tree-based GENIE3 algorithm (Huynh-Thu et al., 2010), enabling the inference of candidate TFs and their regulatory targets (Fig. 1). In parallel, we applied weighted gene correlation network analysis (WGCNA) (Langfelder and Horvath, 2008) to identify stress-specific co-expression modules. Combining these approaches provides complementary insights: while GENIE3 predicts directional regulatory interactions, WGCNA captures correlation-based gene clusters that may reflect shared regulatory control. The overlap between modules identified by both methods suggests that co-expression patterns often arise from underlying TF-driven mechanisms. Furthermore, stress-specific TFs were validated by transcription factor binding site (TFBS) motif enrichment within the promoters of predicted target genes using the Paired Motif Enrichment Tool (PMET, https://pmet.online/) (Rich-Griffin et al., 2020). Overall, this pipeline demonstrates how publicly available transcriptome data can be used to uncover key regulatory mechanisms and generate hypotheses for future research.

**Fig. 1.**
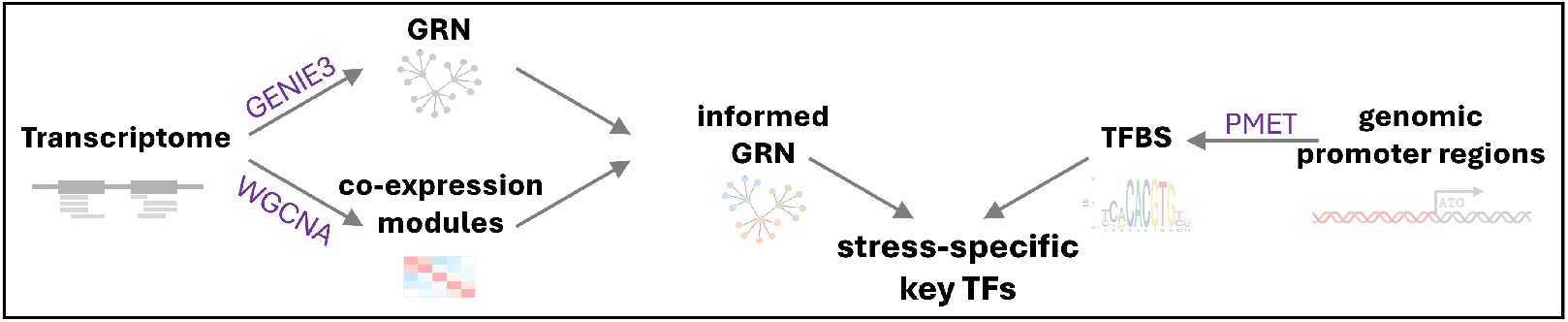
Workflow. Transcriptomic data were analyzed to infer a GRN using GENIE3 and to identify co-expression modules via WGCNA. Integrating both approaches led to an informed GRN that enables the identification of module- and stress-specific key TFs. Promoter regions of predicted target genes were further examined for co-occurring TFBS motifs using PMET. Matching TFBS motifs with corresponding key TFs provides additional support for dataset quality and the robustness of the analysis.

## Materials and Methods

All used scripts are available on GitLab (https://gitlab.lrz.de/christina.steidele/barley-networkinference).

### Overview of samples

Table 1 provides an overview of the experimental design for the 3’RNA-sequencing data deposited on the GEO webpage with the project number GSE223521. Four different barley cultivars (Barke, Grace, Morex, and Palmella Blue) were grown under two treatments: drought stress starting 7 days before infection and normal irrigation. Additionally, three different inoculations were applied: *F. culmorum, F. avenaceum*, and a mock treatment. Samples were collected 48 hours post-infection (hpi) and again at 96 hpi (Hoheneder et al., 2023).

**Table 1.** 3’RNA-sequencing sample overview used in this study and published with the project number GSE22352 on the GEO webpage.

| Cultivar | Treatment | Inoculation | Time point |
| --- | --- | --- | --- |
| Barke | drought | *F. culmorum* | 48hpi |
| Grace |  | *F. avenaceum* |  |
| Morex | irrigated | mock | 96hpi |
| Palmella Blue |  |  |  |
| 3 replicates |  |  |  |

### 3’RNA-sequencing data

#### Mapping

The 3’RNA-sequencing dataset (GSE223521) was mapped against a combined reference comprising the annotated barley genome Morex V3 (Mascher et al., 2021) and the *F. culmorum* UK99 genome (Urban et al., 2016). Mapping and quantification were performed using the nfcore/rnaseq pipeline (version 3.9) (Ewels et al., 2022), employing STAR (Dobin et al., 2013) for read alignment and Salmon (Patro et al., 2017) for transcript quantification. For downstream analyses, only reads mapped to the barley genome were retained.

#### Fungal read counts as external traits for WGCNA

The degree of fungal infection was estimated by calculating the ratio of the total reads mapped to the fungal genome to the total reads mapped to the barley genome. This ratio served as an external trait in the WGCNA analysis.

#### Filtering

The raw barley count table was filtered to retain 20,544 genes with at least 10 reads in a minimum of 10% of samples. Normalization of the filtered counts was performed using DESeq2 (version 1.42.2) (Love et al., 2014), which estimates size factors to correct for differences in sequencing depth. To stabilize variance across the range of mean values, a regularized log (rlog) transformation was applied.

#### Differentially Expressed Genes

Differential expression analysis was conducted using DESeq2 (version 1.42.2) (Love et al., 2014). Treated samples were compared against their respective controls (mock-treated and irrigated). A complete list of differentially expressed genes (DEGs) and their WGCNA module assignments is provided in the supplemental table Supp.Table1.

### Gene regulatory network inference

#### Definition of regulators

The GENIE3 algorithm expects a list of regulators, for which the regulatory weight to all other genes will be calculated. In the absence of an annotated table of barley TFs, a manual search was performed based on the provided genome annotation and included search terms like “transcription factor”, but also well-described gene family names like “WRKY” found on the webpage “Plant transcription factor database” (https://planttfdb.gao-lab.org/) (Jin et al., 2017).

#### Network inference

The gene regulatory network was inferred with the GENIE3 R-package (version 1.24.0) (Huynh-Thu et al., 2010).

#### Louvain clustering

The obtained GRN was filtered for the 10,000 highest edge weights and the Louvain-Clustering algorithm (Blondel et al., 2008) implemented in the igraph R-package (version 2.0)(Csardi and Nepusz, 2006) was used to calculate network clusters based on the GRN adjacency matrix.

### Weighted gene correlation network analyses (WGCNA)

#### Input data

The filtered and normalized count table was used as input for the WGCNA.

#### External trait - Abscisic Acid

Abscisic Acid content was determined using Mass spectrometry according to the methods described by (Chaudhary et al., 2020). 150 mg of ground spike tissue, identical to the material used for 3’RNA-sequencing, was used for analysis. The results are published in Hoheneder et al. (2023).

#### WGCNA

Given the extremely low infection rate of *F. avenaceum* in this experiment, samples inoculated with this species were excluded from the WGCNA. The weighted gene co-expression network analysis (WGCNA) was performed using the WGCNA R package (version 1.73) (Langfelder and Horvath, 2008) with the following parameters: blockwiseModules(myDatExpr, power = 6, TOMType = “signed”, minModuleSize = 30, reassignThreshold = 0, mergeCutHeight = 0.25, numericLabels = TRUE, pamRespectsDendro = FALSE, saveTOMs = TRUE, saveTOMFileBase = “myDatTOM”, verbose = 3). Modules were assigned default color labels by the package. Heatmaps were generated using the pheatmap R package (version 1.0.12) (Raivo, 2019).

### Module-cluster-overlaps and enrichment scores

#### Contingency Table and Enrichment Analysis

To assess the association between Louvain cluster membership and WGCNA module assignment, a contingency table was constructed from the observed counts of genes in each cluster-module combination. Expected counts were calculated under the assumption of independence using the formula:

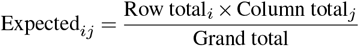

Enrichment was computed as the ratio of observed to expected counts, and log_2_-transformed enrichment values were capped between −5 and 5 for visualization.

#### Filtering and Statistical Testing

For each cluster-module pair, Fisher’s exact test was applied to evaluate whether the observed overlap was greater than expected by chance. To account for multiple comparisons, p-values were adjusted using the Benjamini–Hochberg procedure to control the false discovery rate (FDR) (Benjamini and Hochberg, 2018).

#### Relative weighted node out-degree

The relative weighted node out-degree for TFs was calculated using GENIE3 network inference results combined with WGCNA module assignments. First, edges from the GENIE3 network were filtered by a minimum weight threshold of 0.015 to retain strong regulatory interactions. The edge-weight threshold was determined by k-means clustering on GENIE3 weights, selecting k = 3 via the elbow criterion and using the minimum weight of the high-confidence cluster. For each selected WGCNA module, the module genes were identified and their regulators extracted from the filtered network. Relative weighted node out-degree was calculated as the sum of regulatory weights for all predicted regulators of a specific module and normalized by the number of genes in the module.

#### PMET

Overlapping gene identifiers between Louvain clustering and WGCNA modules were analyzed using PMET (https://pmet.online/) against the precomputed promoter regions of the *Hordeum vulgare* Morex V3 genome. Plant-TFDB (https://planttfdb.gao-lab.org/) (Jin et al., 2017) was used as the motif database, and default parameters were applied (Promoter length: 1000 bp; Max motif matches: 5; Selected promoters: 5000; FIMO threshold: 0.05; Information content threshold: 4; 5UTR included; overlapping promoters removed). Results were filtered for significance using Bonferroni-adjusted p-values (< 0.05).

## Results

### Inference and clustering of a gene regulatory network

GENIE3 (Gene Network Inference with Ensemble of Trees) uses an ensemble of decision trees to infer GRNs from gene expression data (Huynh-Thu et al., 2010). Specifically, the method predicts the expression of each target gene based on the expression of all other genes. The feature importance of each input gene is used as an indicator of a potential regulatory link. We utilized the published 3’RNA-sequencing experiment (Hoheneder et al., 2023) conducted on barley under single (drought or FHB) and double stress (drought and FHB) conditions. To infer the GRN, we used GENIE3 (Huynh-Thu et al., 2010). To reduce noise and focus on the most confident interactions, the top 10,000 edges with the highest regulatory weights were arbitrarily selected and clustered using the Louvain algorithm (Blondel et al., 2008), enabling the identification of gene groups predicted to be co-regulated by the same TFs. These 10,000 edges correspond to a total of 3,748 regulated genes in the GRN. Louvain clustering is a community detection algorithm (Blondel et al., 2008) based on modularity optimization, where it iteratively groups nodes into communities by maximizing the difference between actual and expected edge densities, making it useful for uncovering connectivity patterns within complex networks. The network was plotted, and the 10 biggest Louvain clusters are highlighted with different colors (Fig. 2B).

**Fig. 2.**
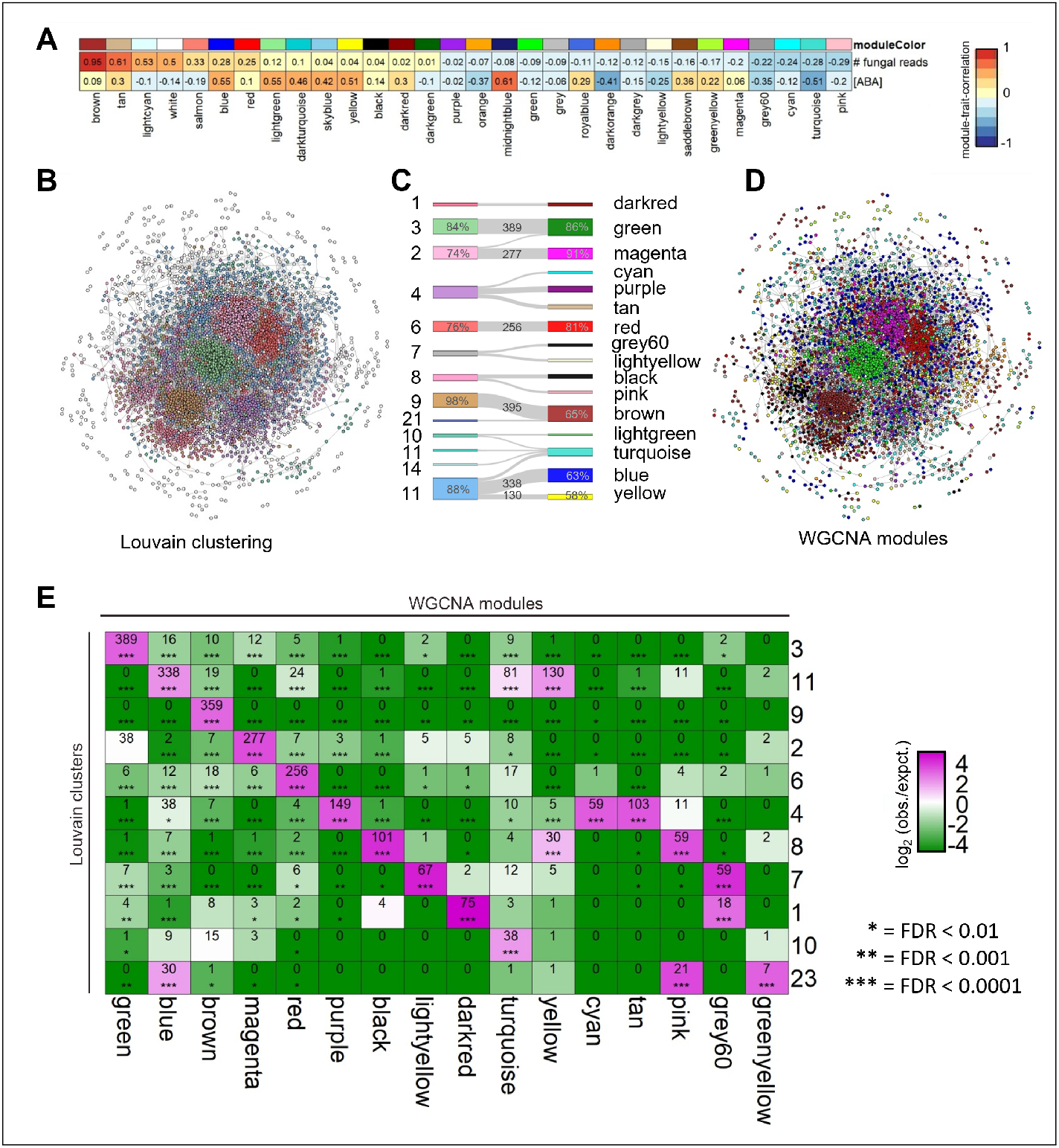
Overlap between Louvain-Clusters and WGCNA modules. **(A)** A WGCNA was performed, resulting in modules linked to fungal read counts as a marker for infection and Abscisic Acid as a marker for drought stress. The correlation of identified co-expression modules with both traits is color-coded, ranging from -1 (blue) to 1 (red). The numbers indicate the module trait correlation. Module names are indicated at the top, with their corresponding names displayed below. **(B)** The 10,000 highest edge weights are plotted, with the 11 largest Louvain clusters color-coded. In the same network layout, the nodes are color-coded by their respective WGCNA module colors in panel **(D)**. Panel **(C)** shows a Sankey diagram illustrating overlaps of more than 30 genes between the largest Louvain clusters and their corresponding WGCNA modules. Grey numbers indicate the number of shared genes between each cluster–module pair. Percentages represent the proportion of shared genes relative to the size of the respective cluster or module. **(E)** Enrichment scores as the log_2_(observed*/*expected) overlaps (green for depletion, pink for enrichment), together with Benjamini-Hochberg corrected FDR values (significance levels are indicated as ∗ = FDR *<* 0.01, ∗∗ = FDR *<* 0.001, ∗ ∗ ∗ = FDR *<* 0.0001) are plotted. The numbers indicate the overlapping genes between a Louvain cluster and a WGCNA module.

#### WGCNA identifies drought- and FHB-associated co– expression modules

WGCNA is an unsupervised approach based on network theory, widely utilized to analyze transcriptomic data and identify co-expressed genes. This method performs pairwise comparisons and allows for the correlation of measured traits with co-expressed genes, helping to associate biological functions (Langfelder and Horvath, 2008). The WGCNA organized the genes into 29 modules of varying sizes and labels them by different random colors. Module sizes range from 3904 genes (turquoise module) to 48 genes (saddlebrown module), and one cluster with 1359 genes where all remaining genes were collected (grey module) (Fig. 2A). The brown module, consisting of 2,950 genes, shows a significant correlation of 0.95 with the fungal read RNA count, which is the highest correlation observed in this analysis (Fig. 2A). The fungal read counts in the 3’RNA-sequencing data are a marker for FHB severity; higher fungal read counts indicate greater fungal biomass and, consequently, a more severe infection. Therefore, we will refer to the brown module as the infection-specific module. For the concentration of the drought stress hormone abscisic acid (ABA) measured in parallel from the same samples, the midnightblue module (188 genes) exhibits the highest correlation at 0.61, followed closely by the blue module (3,262 genes) and the lightgreen module (167 genes) with a correlation of 0.55. The yellow module (1,633 genes) has a correlation of 0.51 with ABA. We refer to the two largest ABA-correlated modules (blue and yellow) as drought stress modules. The WGCNA proved to be very valuable for this dataset, because it helped to reduce over 20,500 genes to around 2600 that are likely important during FHB response and roughly 4,800 genes involved in the drought stress response by identifying the largest modules, positively associated with the given trait.

#### Co-expressed genes cluster within the GRN

To locate the genes associated with drought or infection within the GRN, the nodes were color-coded according to their WGCNA module membership (Fig. 2A) using the same filtering and network layout as in Fig. 2B (Fig. 2D). This illustrates large overlaps of GRN clusters and WGCNA coexpression modules, suggesting that co-expression patterns can be traced back to TF-driven gene regulatory networks. To further evaluate the relationship between Louvain clusters and WGCNA modules, we used a Sankey diagram to visualize overlaps between the two different clustering approaches, applying a minimum threshold of 30 shared genes between cluster-module pairs (Fig. 2C). We identified several module-cluster pairs with significantly enriched overlaps (Fig. 2E). For instance, 359 genes from the brown infection-specific module (65% of that module) overlap with Louvain cluster 9 (98% of that cluster). Cluster 11 (626 genes) overlaps mainly with the blue and yellow modules: 338 genes belong to the blue module (63% of blue; 54% of cluster 11) and 130 genes to the yellow module (58% of yellow; 21% of cluster 11). By contrast, the majority of cluster-module combinations display negative log_2_-(observed/expected) enrichment score values, suggesting that the observed overlap is lower than expected under a random distribution of genes. The comparison between the results of the tree-based GENIE3 algorithm, which infers regulatory interactions, with the pairwise correlation-based WGCNA approach suggests that co-expression patterns may result from underlying common regulatory mechanisms driven by TFs. Despite their different mathematical frameworks, both methods yielded strongly overlapping gene modules of probable biological relevance.

#### Drought Stress and FHB response are regulated by module-specific key transcription factors

To identify key TFs potentially regulating the response to drought stress or FHB, we calculated the relative weighted out-degree for all predicted regulators within the blue and yellow modules (associated with drought stress) and the brown module (associated with infection) (Fig. 3A). We included the green and red cluster here as clusters exhibiting the largest overlaps between Louvain-clustering and WGCNA co-expression modules, which, however, lack a clear physiological association with any of the applied stresses. Our analysis revealed minimal overlap among the TFs with the highest relative weighted out-degree in each module, indicating that each module is regulated by a distinct set of key TFs. These key TFs represent regulators with the strongest predicted influence on target genes within their respective modules. Only the drought stress-associated blue and yellow modules shared a subset of such TFs with relatively high out-degree values, which is expected as they are grouped together in Louvain cluster 11 (Fig. 2C).

**Fig. 3.**
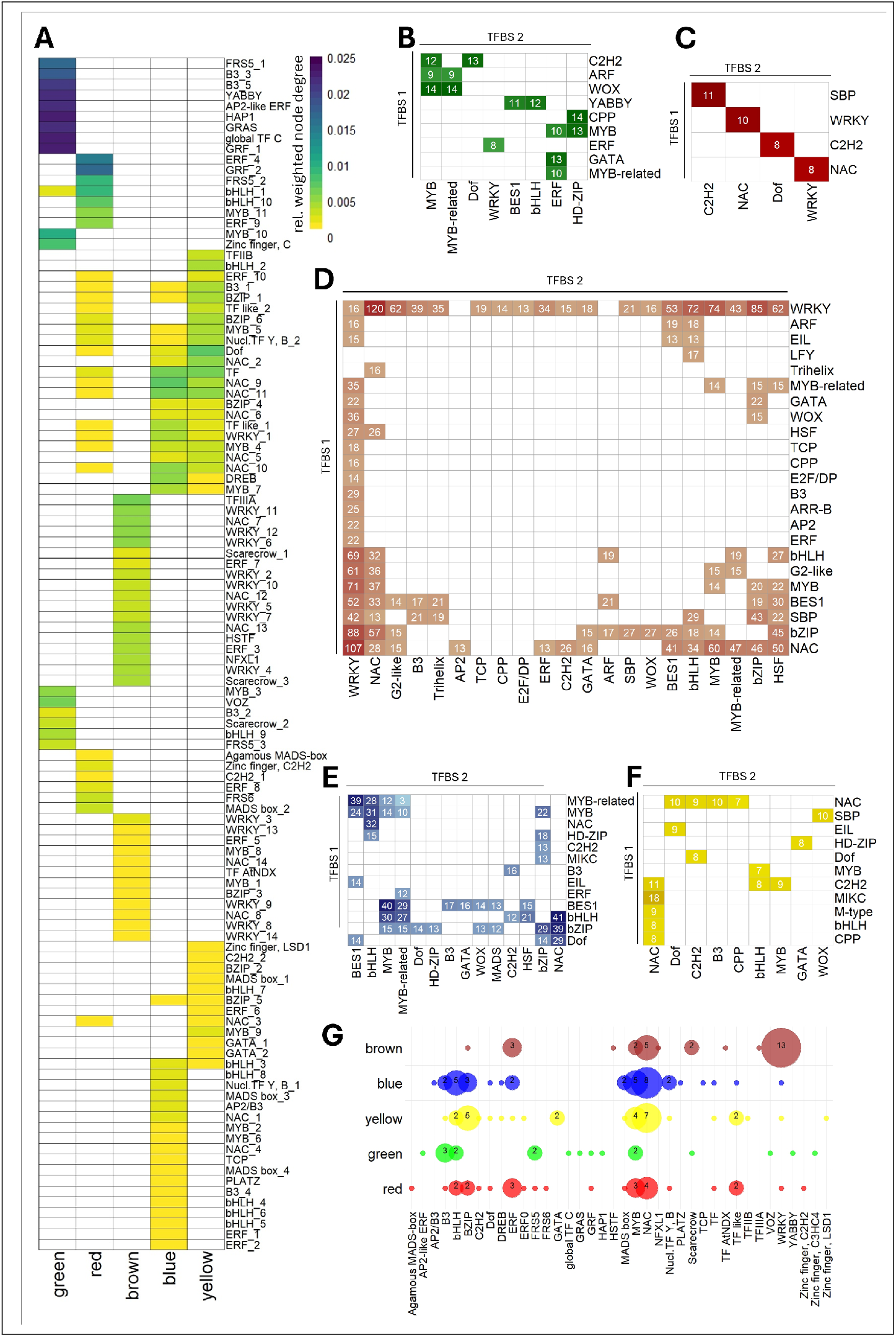
Link between key TFs and TFBSs occurrence across selected modules. **(A)** The relative weighted node out-degree for all predicted regulators of the green, red, brown, yellow and blue modules were calculated and all values > 0.001 are shown. PMET was used to identify significantly enriched TFBS motifs in the promoter regions of the genes (Bonferroni p-value < 0.05) within the overlaps of **(B)** green/3, **(C)** red/6, **(D)** brown/9, **(E)** blue/11 and **(F)** yellow/11 module/cluster. The numbers indicate the number of identified promoters carrying the respective motif pair based on genes that overlapped between GENIE3 and WGCNA analyses. **(G)** shows the numbers of key TFs, ordered by families, derived from A.

#### Module-dependent enrichment of TFBSs

To determine whether the identified key TFs could potentially regulate the expression of their inferred target genes, we screened their 1 kb upstream promoter regions for the presence of co-occurring known transcription factor binding site (TFBS) motifs using the Paired Motif Enrichment Tool (PMET, ) (Rich-Griffin et al., 2020). We focused on genes that overlapped between the GRN inferred by GENIE3 and the co-expression modules identified by WGCNA, as these genes were detected through two independent approaches (Fig. 2E). The following numbers of genes shared between WGCNA modules and GENIE3 clusters were obtained: 209 genes in the green module and cluster 3, 152 genes in the red module and cluster 6, 320 genes in the brown module and cluster 9, 306 genes in the blue module and cluster 11, and 117 genes in the yellow module and cluster 11 (Fig. 2E).

We identified significantly enriched pairs of TFBS motifs for each set of clustered genes (Bonferroni-adjusted *p*-value < 0.05). In the green module, MYELOBLASTOSIS (MYB) and MYB-like TFBSs frequently co-occurred with WUSCHEL-related homeobox (WOX) motifs (Fig. 3B). In the red module, 11 genes contained a Cys2-His2 zinc finger protein (C2H2) motif together with a SQUAMOSA Promoter Binding Protein (SBP) motif (Fig. 3C). The brown, FHB-related module was dominated by WRKY motifs strongly associated with NAC TFBSs (WRKY–NAC: 107 genes, 33%; NAC–WRKY: 120 genes, 37.5%), but also showed co-occurrence with several other TFBSs (Fig. 3D). Promoter regions of the blue, drought-stress module were enriched for NAC motifs, which, in contrast to the brown infection module, co-occurred with basic helix–loop–helix (bHLH) motifs. Additionally, the blue module showed strong enrichment for BRASSINOSTEROID INSENSITIVE1–EMS–SUPPRESSOR1 (BES1)/MYB motif pairs (Fig. 3E), whereas in the brown module BES1 was associated with NAC, WRKY, and Heat Shock Transcription Factor (HSF) motifs, but not with MYB-related motifs. In the yellow, drought-stress module, 18 promoters contained a NAC motif together with a MADS-box transcription factor of the MIKC type (MIKC) binding motif, and 11 promoters showed NAC motifs co-occurring with C2H2 motifs (Fig. 3F), both combinations are absent from the blue module. The distinct enrichment patterns of TFBS motif pairs across modules reflect the unique co-expression signatures identified by WGCNA and their corresponding regulatory clusters in the gene regulatory network, suggesting module-specific transcriptional regulation.

#### TFBS motifs align with inferred key TFs

We performed a comparative analysis of the families of key TFs from each module, examining their co-occurring TFBSs in putative target genes. For the green module, we identified 12 YABBY motifs associated with the BES1 TFBS motif and 11 motifs linked to bHLH (Fig. 3B). Notably, the only YABBY domain-containing (YABBY) TF identified among all key TFs is specifically grouped to regulate the green module (see Fig. 3G). In contrast, the genes within the FHB-responsive, brown module show significant enrichment for promoters featuring WRKY TFBSs, with 13 out of 16 predicted key WRKY TFs being strongly associated with this module (Fig. 3G). The BES1 motif is also present in the brown module and is significantly enriched alongside other motifs such as NAC, basic leucine zipper (bZIP), WRKY, and HSF (Fig. 3D). However, it is not associated with MYB and MYB-like TFBSs, as seen in the blue module (Louvain cluster 3; Fig. 3E). The drought-responsive, blue and the yellow module are both significantly associated with ABA (Fig. 2A) and are partially clustered together in the GRN in the Louvain cluster 11 (Fig. 2C). They share a set of overlapping key TFs but are predicted to be regulated by also distinct sets of key TFs (Fig. 3A). In both modules, NAC TFBS motifs frequently cooccurred with other motifs. Among the three key transcription factors with the highest relative weighted node degrees in the blue and yellow modules, two are NAC TFs (Fig. 3A). Within the distinct key TFs of the yellow module, there is a C2H2 TF and a MADS-box TF (Fig. 3A), aligning with the predicted C2H2 and MIKC motif (Fig. 3F). Taken together, these findings underscore the consistency between motif enrichment patterns and the predicted key transcription factors, supporting the relevance of the identified key TFs within each module, previously identified based on regulatory weight in transcriptomic data.

## Discussion

We developed a bioinformatics pipeline (Fig. 1) for reanalyzing the publicly available 3RNA-sequencing dataset from a barley greenhouse experiment involving combined FHB and drought stress to investigate transcriptional regulation in stress-responsive genes. To uncover regulatory patterns, we applied two complementary mathematical frameworks: WGCNA, which identifies co-expressed gene modules through correlation-based hierarchical clustering, and GENIE3, which infers directional GRNs using machine learning. This analysis was further complemented by independent identification of TFBS motif pairs in the promoter regions of putative TF target genes.

WGCNA (Langfelder and Horvath, 2008) was employed to identify modules of co-expressed genes associated with drought stress or FHB response. This analysis revealed a major module (brown module) strongly associated with the number of fungal reads detected in each sample (Figure 2A). The 3’RNA-sequencing dataset was originally published by Hoheneder et al. (2023), where co-expression modules were correlated with quantified fungal DNA from pooled spike material. In contrast, our analysis achieved a substantially higher correlation (0.95 vs. 0.73) because it was performed on read data from individual samples rather than fungal DNA from pooled samples. The number of fungal reads can be considered an *in silico* proxy for total fungal gene transcripts and serves as a robust marker for fungal infection.

Using GENIE3, we constructed a GRN to predict regulatory interactions between TFs and all expressed genes based on transcriptomic profiles. The resulting network was partitioned into clusters using Louvain clustering (Blondel et al., 2008). We integrated the results from both approaches to identify gene-sets being predicted to be both co-expressed and regulated by the same TFs. The WGCNA identified co-expressed genes significantly associated with either drought stress (blue and yellow module) or FHB response (brown module). The Louvain clustering algorithm partitions the GRN into modules that are more densely interconnected internally than with the rest of the network. Since the GENIE3-based GRN is a directed network, with edges pointing from TFs to their target genes, each Louvain module tends to be organized around a distinct set of TFs with overlapping target genes. The significant overlaps between the drought stress-specific blue and yellow modules with Louvain cluster 11, as well as the overlap between the FHB-responsive brown module and cluster 9, suggest that the co-expression of these genes may be regulated by a unique, module-specific set of key TFs. Our analysis revealed both statistically significant overlaps and cases where the observed overlap between Louvain clusters and WGCNA modules fell below random expectation. This consistency across two distinct mathematical frameworks — network-based clustering and co-expression analysis — suggests a strong biological difference between sample groups. While such an agreement is theoretically expected, it is not always observed in biological data, underscoring the robustness of our approach and the dataset used. To identify key TFs influencing drought or FHB response, we calculated the relative weighted node out-degree for all TFs within the brown, blue, yellow, red, and green modules. TFs with the highest weighted node-degrees are predicted to have the strongest influence on the expression patterns of their respective modules. Each module is regulated by a distinct set of key TFs, as suggested by the overlap of Louvain and WGCNA clusters. Interestingly, these TF sets differ greatly in their family composition, with MYB TFs being the only family present in all five evaluated modules, indicating highly tailored gene expression regulation. This corroborates our earlier assumption that the barley stress response reflects a modular composition of separated biotic and abiotic stress responses that modularly contribute to a combined stress response with few gene expression patterns that cannot be explained from additive single stress responses (Hoheneder et al., 2023).

To further validate our findings, we employed an independent approach using PMET (Rich-Griffin et al., 2020), which identifies significantly enriched pairs of TFBS motifs within promoter regions. PMET focuses on motif pairs because combinatorial regulation by multiple TFs is a key mechanism in complex stress responses. This analysis revealed enriched cis-element pairs in promoters of co-expressed genes that corresponded well to binding sites of predicted TFs. For example, in the infection-specific brown module, key TFs included 13 WRKY family members—a plant-specific TF family widely recognized for roles in biotic and abiotic stress responses (Mahiwal et al., 2024). Among these, HvWRKY23 (HORVU.MOREX.r3.7HG0743280) has previously been shown to modulate defense responses and enhance FHB resistance in barley (Karre et al., 2019). Transient silencing of HvWRKY23 reduced expression of flavonoid and hydroxycinnamic acid amide (HCAA) biosynthetic genes, leading to decreased accumulation of resistance-related metabolites and increased susceptibility. Promoter-binding luciferase assays confirmed that HvWRKY23 activates promoters of several flavonoid glucoside biosynthesis genes (Karre et al., 2019). Recent multi-omics evidence further supports the upregulation of HCAA biosynthetic pathways under stress conditions (Hein et al., 2025). In line with these findings, we identified two tryptophan decarboxylases predicted to be regulated by HvWRKY23 and carrying WRKY TFBS motifs. WRKY family members have been broadly implicated in flavonoid and HCAA biosynthesis across species, including AtWRKY23 in Arabidopsis (auxin-dependent flavonol biosynthesis) (Grunewald et al., 2008), StWRKY1 in potato, and TaWRKY70 in wheat (Yogendra et al., 2015; Kage et al., 2017).

BES1 is a major TF in brassinosteroid (BR)-dependent signaling and is regulated through phosphorylation (Yin et al., 2002, 2005). In the absence of BR, BES1 phosphorylation prevents DNA binding and dimerization with other TFs (Han et al., 2023). BR signaling generally promotes growth, whereas stress-induced ABA signaling is typically antagonistic to BR signaling. In the blue module, BES1 TFBS motifs co-occur with MYB and MYB-related motifs (Fig. 3E). The MYB TF family is one of the largest in plants, regulating processes such as stress tolerance, development, and secondary metabolite biosynthesis. In contrast, in the brown module, BES1 motifs co-occur with NAC, WRKY, or HSF motifs (Fig. 3D), indicating a more stress-responsive regulatory network. This pattern suggests a mechanism for finetuning the growth–immunity trade-off: BES1 may integrate BR-driven growth signals with stress-related transcriptional programs by partnering with different TF families depending on physiological context.

These observations provide additional support for the inferred GRN and strengthen a putative role of the suggested key TFs as drivers of stress-specific responses. However, further analysis is needed to assign individual TFBS motifs to specific TF family members and validate these predictions *in planta*. Additionally, most predicted TFBS motifs are derived from other plant species and should be carefully evaluated for barley. This is particularly relevant because TF families differ in motif conservation: NAC, WRKY, BES1, and MADS-MIKC members bind highly similar consensus motifs, whereas families such as bHLH, bZIP, MYB, and MYB-related exhibit only semi-conserved motifs. In contrast, AP2/EREB, C2H2, C3H, and Trihelix families bind to a broader variety of motifs (Zenker et al., 2025). These differences highlight the complexity of motif-based predictions and the need for species-specific validation.

In summary, we employed two complementary methods to identify co-regulated gene sets using WGCNA and GENIE3. Both approaches aim to uncover relationships in gene expression, but rely on different mathematical principles. Our analyses were based on transcriptomic data, and the significantly overlapping results were further supported by analysis of genomic TFBSs within the promoter regions of TF target genes, using PMET. This allowed us to identify key TFs that may drive the response to drought stress and FHB. Additionally, we confirmed that the putative target genes contain TFBSs corresponding to these predicted key TFs. Data further support largely separated gene regulatory mechanisms for abiotic and biotic stress responses in barley.

## Supporting information

Supplemental Table 1 - DEGs

## ACKNOWLEDGEMENTS

We thank Prof. Dr. Patrick Schäfer for fruitful discussions and his suggestion to use PMET.

## Bibliography

Y. Benjamini and Y. Hochberg. Controlling the false discovery rate: A practical and powerful approach to multiple testing. Journal of the Royal Statistical Society: Series B (Methodological), 57, 2018.

V.D. Blondel, J.L. Guillaume, and E. Lambiotte, R. and Lefebvre. Fast unfolding of communities in large networks. Journal of Statistical Mechanics: Theory and Experiment, 2008, 2008.

H. Buerstmayr and M. Lemmens. Breeding healthy cereals: genetic improvement of fusarium resistance and consequences for mycotoxins. World Mycotoxin Journal, 8, 2015.

A. Chaudhary, X. Chen, J. Gao, B. Leśniewska, R. Hammerl, C. Dawid, and K. Schneitz. The arabidopsis receptor kinase strubbelig regulates the response to cellulose deficiency. PLOS Genetics, 16, 2020.

G. Csardi and T. Nepusz. The igraph software package for complex network research. InterJournal, Complex Systems, 1695, 2006.

M. Delgado-Baquerizo, C.A. Guerra, C. Cano-Díaz, E. Egidi, J. Wang, N. Eisenhauer, B.K. Singh, and F.T. Maestre. The proportion of soil-borne pathogens increases with warming at the global scale. Nature Climate Change, 10, 2020.

A. Dobin, C.A. Davis, F. Schlesinger, J. Drenkow, C. Zaleski, S. Jha, P Batut, M. Chaisson, and T.R. Gingeras. Star: ultrafast universal rna-seq aligner. Bioinformatics, 29, 2013.

P. Ewels, A. Peltzer, S. Fillinger, H. Patel, J. Alneberg, A. Wilm, M. Ulysse Garcia, P. Di Tommaso, and S. Nahnsen. The nf-core framework for community-curated bioinformatics pipelines., 2022.

N.A. Foroud, D. Baines, T.Y. Gagkaeva, N. Thakor, A. Badea, B. Steiner, M. Bürstmayr, and H. Bürstmayr. Trichothecenes in cereal grains – an update. Toxins, 11, 2017.

W. Grunewald, M. Karimi, K. Wieczorek, E. Van de Cappelle, E. Wischnitzki, F. Grundler, D. Inzé, T. Beeckman, and G. Gheysen. A role for atwrky23 in feeding site establishment of plantparasitic nematodes. Plant Physiology, 148, 2008.

C. Han, L. Wang, J. Lyu, W. Shi, L. Yao, M. Fan, and M.-Y. Bai. Brassinosteroid signaling and molecular crosstalk with nutrients in plants. Journal of Genetics and Genomics, 50, 2023.

S. Hein, C.E. Steidele, F. Hoheneder, S. Brajkovic, B. Kuster, L. Kurzweil, T.D. Stark, C. Dawid, and R. Hückelhoven. Multi-omics of barley fusarium head blight converge on pathogen-triggered biosynthesis of aromatic amino acid derived chemical defense compounds. bioRxiv, 2025.

F. Hoheneder, C.E. Steidele, M. Messerer, K.F.X. Mayer, N. Köhler, C. Wurmser, M. Heß, M. Gigl, C. Dawid, R. Stam, and R. Hückelhoven. Barley shows reduced fusarium head blight under drought and modular expression of differentially expressed genes under combined stress. Journal of Experimental Botany, 74, 2023.

F. Hoheneder, C.E. Steidele, M. Gigl, C. Dawid, and R. Hückelhoven. Transcriptome and hormone regulations shape drought stress-dependent fusarium head blight susceptibility in different barley genotypes. bioRxiv, 2025.

Y. Huang, L. Yin, A.H. Sallam, S. Heinen, L. Li, K. Beaubien, R. Dill-Macky, Y. Dong, B.J. Steffenson, K.P. Smith, and G.J Muehlbauer. Genetic dissection of a pericentromeric region of barley chromosome 6h associated with fusarium head blight resistance, grain protein content and agronomic traits. Theoretical and Applied Genetics, 134, 2021.

V.A. Huynh-Thu, A. Irrthum, and P. Wehenkel, L. and Geurts. Inferring regulatory networks from expression data using tree-based methods. PLoS ONE, 5, 2010.

J.P. Jin, F. Tian, D.C. Yang, Y.Q. Meng, L. Kong, J.C. Luo, and G. Gao. Planttfdb 4.0: toward a central hub for transcription factors and regulatory interactions in plants. Nucleic Acids Research, 45, 2017.

U. Kage, K.N. Yogendra, and A.C. Kushalappa. Tawrky70 transcription factor in wheat qtl-2dl regulates downstream metabolite biosynthetic genes to resist fusarium graminearum infection spread within spike. Scientific Reports, 7, 2017.

S. Karre, A. Kumar, K. Yogendra, U. Kage, A. Kushalappa, and J.B. Charron. Hvwrky23 regulates flavonoid glycoside and hydroxycinnamic acid amide biosynthetic genes in barley to combat fusarium head blight. Plant Molecular Biology, 100, 2019.

A. Kumar, K.N. Yogendra, S. Karre, A.C. Kushalappa, Y. Dion, and T.M. Choo. Wax inducer1 (hvwin1) transcription factor regulates free fatty acid biosynthetic genes to reinforce cuticle to resist fusarium head blight in barley spikelets. Journal of Experimental Botany, 67, 2016.

P. Langfelder and S. Horvath. Wgcna: an r package for weighted correlation network analysis. BMC Bioinformatics, 9, 2008.

H. Li, J. Wu, X. Shang, M. Geng, J. Gao, S. Zhao, X. Yu, D. Liu, Z. Kang, X. Wang, and X. Wang. Wrky transcription factors shared by bth-induced resistance and npr1-mediated acquired resistance improve broad-spectrum disease resistance in wheat. Molecular Plant-Microbe Interactions®, 33, 2020.

A. Linkmeyer, M. Götz, L. Hu, S. Asam, M. Rychlik, H. Hausladen, M. Hess, and R. Hückelhoven. Assessment and introduction of quantitative resistance to fusarium head blight in elite spring barley. Phytopathology, 104, 2013.

M.I. Love, W. Huber, and S. Anders. Moderated estimation of fold change and dispersion for rna-seq data with deseq2. Genome Biology, 15, 2014.

S. Mahiwal, S. Pahuja, and G.K. Pandey. Review: Structural-functional relationship of wrky transcription factors: Unfolding the role of wrky in plants. International Journal of Biological Macromolecules, 257, 2024.

M. Mascher, T. Wicker, J. Jenkins, C. Plott, T. Lux, C.S. Koh, J. Ens, H. Gundlach, L.B. Boston, Z. Tulpová, S. Holden, I. Hernández-Pinzón, U. Scholz, K.F.X. Mayer, M. Spannagl, C.J. Pozniak, A.G. Sharpe, H. Šimková, M.J. Moscou, J. Grimwood, J. Schmutz, and N. Stein. Longread sequence assembly: a technical evaluation in barley. The Plant Cell, 33, 2021.

T. Miedaner, C. Flamm, and M. Oberforster. The importance of fusarium head blight resistance in the cereal breeding industry: Case studies from germany and austria. Plant Breeding, 143, 2024.

P. Ogrodowicz, A. Kuczyńska, K. Mikołajczak, T. Adamski, M. Surma, P. Krajewski, H. ćwiek Kupczyńska, M. Kempa, M. Rokicki, and D. Jasińska. Mapping of quantitative trait loci for traits linked to fusarium head blight in barley. PLOS ONE, 15, 2020.

P. Pandey and M. Senthil-Kumar. Plant-pathogen interaction in the presence of abiotic stress: What do we know about plant responses? Plant Physiology Reports, 24, 2019.

D.W. Parry, P. Jenkinson, and McLeod L. Fusarium ear blight (scab) in small grain cereals—a review. Plant Pathology, 44, 1995.

R. Patro, G. Duggal, M.I. Love, R.A. Irizarry, and C. Kingsford. Salmon provides fast and biasaware quantification of transcript expression. Nature Methods, 14, 2017.

K. Raivo. pheatmap: Pretty Heatmaps, 2019. URL https://CRAN.R-project.org/package=pheatmap. R package version 1.0.12.

C. Rich-Griffin, R. Eichmann, M.U. Reitz, S. Hermann, K. Woolley-Allen, P.E. Brown, K. Wiwatdirekkul, E. Esteban, A. Pasha, K.-H. Kogel, N.J. Provart, S. Ott, and P. Schäfer. Regulation of cell type-specific immunity networks in arabidopsis roots. The Plant Cell, 32, 2020.

A.H. Sallam, M. Haas, Y. Huang, Z. Tandukar, G. Muehlbauer, K.P. Smith, and B.J. Steffenson. Meta-analysis of the genetics of resistance to fusarium head blight and deoxynivalenol accumulation in barley and considerations for breeding. Plant Breeding, 143, 2024.

M. Urban, R. King, A. Andongabo, U. Maheswari, H. Pedro, P. Kersey, and K. Hammond-Kosack. First draft genome sequence of a uk strain (uk99) of fusarium culmorum. Genome Announcements, 4, 2016.

M. Vranić, A. Perochon, and F.M. Doohan. Transcriptional profiling reveals the wheat defences against fusarium head blight disease regulated by a nac transcription factor. Plants, 12, 2023.

W. Xie, W. Xiong, J. Pan, T. Ali, Q. Cui, D. Guan, J. Meng, N.D. Mueller, E. Lin, and Davis. S.J. Decreases in global beer supply due to extreme drought and heat. Nature Plants, 4, 2018.

Y. Yin, Z.-Y. Wang, S. Mora-Garcia, J. Li, S. Yoshida, T. Asami, and J. Chory. Bes1 accumulates in the nucleus in response to brassinosteroids to regulate gene expression and promote stem elongation. Cell, 109, 2002.

Y. Yin, D. Vafeados, Y. Tao, S. Yoshida, T. Asami, and J. Chory. A new class of transcription factors mediates brassinosteroid-regulated gene expression in arabidopsis. Cell, 120, 2005.

K.N. Yogendra, A. Kumar, K. Sarkar, Y. Li, D. Pushpa, K.A. Mosa, R. Duggavathi, and A.C. Kushalappa. Transcription factor stwrky1 regulates phenylpropanoid metabolites conferring late blight resistance in potato. Journal of Experimental Botany, 66, 2015.

S. Zenker, D. Wulf, A. Meierhenrich, P. Viehöver, S. Becker, M. Eisenhut, R. Stracke, B. Weisshaar, and A. Bräutigam. Many transcription factor families have evolutionarily conserved binding motifs in plants. Plant Physiology, 198, 2025.

